# Eps8 is a convergence point integrating EGFR and integrin trafficking and crosstalk

**DOI:** 10.1101/405043

**Authors:** Nikki R Paul, Joanna R Thomas, Horacio Maldonado, Katarzyna I Wolanska, Ewa J Koper, Jonathan D Humphries, Adam Byron, Adams George, Nathan Allen, Ian A Prior, Charles H Streuli, Martin J Humphries, Mark R Morgan

## Abstract

Crosstalk between adhesion and growth factor receptors plays a critical role in tissue morphogenesis and repair, and aberrations contribute substantially to neoplastic disease. However, the mechanisms by which adhesion and growth factor receptor signalling are integrated, spatially and temporally, are unclear.

We used adhesion complex enrichment coupled with quantitative proteomic analysis to identify rapid changes to adhesion complex composition and signalling following growth factor stimulation. Bioinformatic network and ontological analyses revealed a substantial decrease in the abundance of adhesion regulatory proteins and co-ordinators of endocytosis within 5 minutes of EGF stimulation. Together these data suggested a mechanism of EGF-induced receptor endocytosis and adhesion complex turnover.

Combinatorial interrogation of the networks allowed a global and dynamic view of adhesion and growth factor receptor crosstalk to be assembled. By interrogating network topology we identified Eps8 as a putative node integrating α5β1 integrin and EGFR functions. Importantly, EGF stimulation promoted internalisation of both α5β1 and EGFR. However, perturbation of Eps8 increased constitutive internalisation of α5β1 and EGFR; suggesting that Eps8 constrains α5β1 and EGFR endocytosis in the absence of EGF stimulation. Consistent with this, Eps8 regulated Rab5 activity and was required for maintenance of adhesion complex organisation and for EGF-dependent adhesion complex disassembly. Thus, by co-ordinating α5β1 and EGFR trafficking mechanisms, Eps8 is able to control adhesion receptor and growth factor receptor bioavailability and cellular contractility.

We propose that during tissue morphogenesis and repair, Eps8 functions to spatially and temporally constrain endocytosis, and engagement, of α5β1 and EGFR in order to precisely co-ordinate adhesion disassembly, cytoskeletal dynamics and cell migration.

## INTRODUCTION

Cellular recognition of fibrillar extracellular matrix (ECM) molecules and soluble growth factors leads to localised compartmentalisation of cytoplasmic signalling and topological sensation of the extracellular microenvironment. Integrins are transmembrane adhesion receptors that transmit signals bidirectionally from ECM to the cytoskeleton. Sites of integrin-mediated ECM engagement serve as mechanochemical signalling nexuses integrating the extracellular microenvironment with the contractile machinery of the cell. Integrin-associated complexes (IACs) transmit mechanical forces bidirectionally across the membrane and act as signalling platforms to propagate membrane-distal signals. Thus, regulation of the dynamics of cell-matrix interactions, mechanosensation and mechanotransduction controls cell migration, microenvironment remodelling and global cell fate decisions^1^. However, crosstalk exists between integrins and growth factor receptors (GFRs), which spatially and temporally co-ordinates the functions of both receptor families.

Integrin-GFR crosstalk affects downstream processes such as migration, proliferation and apoptosis^2^; key cell fate decisions that contribute to the maintenance of health and development of disease. Consequently, crosstalk between adhesion and growth factor receptors plays a critical role in tissue morphogenesis and repair, and aberrations contribute substantially to neoplastic progression and therapeutic response^2-4^. Despite this, little is known about the mechanisms that temporally and spatially orchestrate adhesion and GFR crosstalk.

Integrins provide positional cues and enable cells to respond to growth factor signals from the microenvironment, but the role of adhesion receptors is not passive^5^. Integrin-GFR crosstalk can be mediated by a diverse range of mechanisms affecting receptor expression, activity, signalling and trafficking^6^. Thus, adhesion receptor and GFR crosstalk provides mechanisms by which IACs, representing localised foci of mechanical and biochemical signal transduction, are co-ordinated spatially and temporally by GFRs. Equally, integrins can directly influence the subcellular distribution clustering and expression of GFRs^6^. In order to understand how integrins and GFRs regulate biological functions in complex microenvironments, it is necessary to study the complex relationship between the receptor families. However, until recently, it has not been possible to employ global proteomic analyses to dissect the complexity of integrin-GFR crosstalk mechanisms.

IACs comprise large stratified multi-molecular complexes recruited to the cytoplasmic domain of integrins and the integrin-proximal membrane environment, termed the integrin ‘adhesome’. Advances in adhesion isolation techniques and proteomics have enabled unbiased global analysis of adhesion signalling networks and revealed complexity that far exceeds candidate-driven approaches^1, 7-13^. These studies led to identification of thousands of integrin-associated proteins, compared with previous literature curated databases that estimated over 200 components^7, 14-16^. Meta-analysis of IAC proteomic datasets has provided further insight into the complexity of integrin-dependent signalling; leading to the definition of the consensus adhesome and meta-adhesome. The consensus adhesome represents 60 proteins that are consistently recruited to IACs and likely represent the core adhesion machinery. Whereas the meta-adhesome comprises 2,352 proteins that are more variably detected in IACs and may represent proteins that are cell or ECM context-dependent, highly dynamic, of low stoichiometry or labile^7^.

In this study, we employed proteomic analysis of signalling networks established at sites of cell-matrix interaction to identify mechanisms co-ordinating integrin-GFR crosstalk, and to determine the impact of integrin-GFR crosstalk on IAC composition. Network and ontological analysis of isolated IACs, following acute EGF stimulation, revealed an EGF-dependent decrease in adhesion regulatory proteins and co-ordinators of endocytosis, suggesting a mechanism of EGF-induced receptor endocytosis and adhesion complex turnover. Further analyses identified Eps8 as a key regulatory protein integrating α5β1 integrin and EGFR functions. We show that EGF stimulation promotes internalisation of both α5β1 and EGFR and that Eps8 constrains α5β1 and EGFR endocytosis in the absence of EGF stimulation. Consequently, Eps8 is required for maintenance of adhesion complex organisation and for EGF-dependent adhesion complex disassembly. We further show that EGF stimulation activates the endocytic trafficking regulator, Rab5, and that Eps8 controls Rab5 activity. Thus, by co-ordinating α5β1 and EGFR trafficking mechanisms, Eps8 is able to control adhesion receptor and growth factor receptor bioavailability and function.

## RESULTS AND DISCUSSION

### Global analysis of integrin-EGFR crosstalk

To determine the impact of acute EGF stimulation on adhesion signalling, we have established IAC enrichment techniques, coupled with quantitative proteomic analysis, to identify rapid changes to adhesion complex composition following growth factor stimulation. Analysis of MAPK phosphorylation revealed that stimulation of mammary epithelial cells with EGF induced MAPK activation within 5 minutes (Figure 1A). Therefore 5 minutes acute EGF stimulation was selected as the timepoint for integrin-associated complex (IAC) enrichment. IACs were isolated from mammary epithelial cells using hypotonic water pressure following 5 minutes incubation with EGF or serum-free medium. Samples were validated by immunoblotting to ensure specificity of enrichment; i.e. isolation of β1 integrin, paxillin and phospho-Y416 Src, and absence of non-adhesion-specific proteins such as mtHSP70 (Figure 1B). Consistent with the notion that IAC isolation approaches enrich for both "intrinsic" and "associated" components of IACs ^7, 16^, it is notable that active MAPK (pMAPK T202/Y204) was detected in isolated IACs following EGF stimulation.

**Figure 1:**
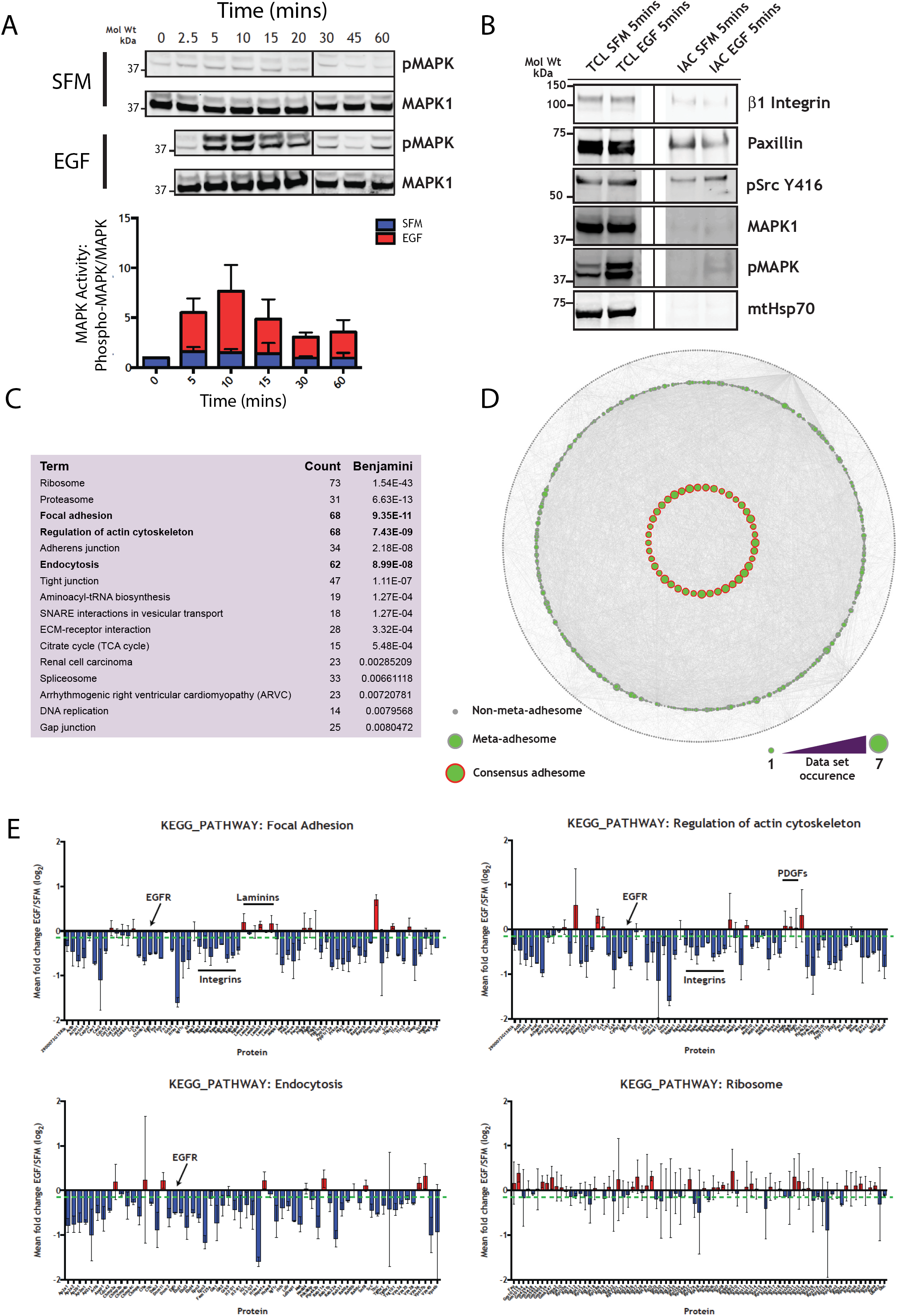
Proteomic and ontological analysis of IACs reveals decreased adhesion regulatory proteins and co-ordinators of endocytosis following EGF stimulation. **A)** MAPK phosphorylation response of MECs following acute EGF treatment. Primary MECs were seeded on collagen-I-coated plates, serum-starved for 4 hours and stimulated with 5ng/ml EGF or maintained in serum-free medium (SFM). Total cell lysates from each timepoint were analysed by immunoblot for total MAPK1 and phospho-MAPK (pMAPK) Quantification of pMAPK1/2 T202/Y204 relative to total MAPK protein levels. N=3, error bars = SEM. Blue bars = Vehicle control, Red bars = EGF stimulation. α-tubulin was used as a loading control (not shown). **B)** Validation of IAC enrichment from EpH4 MECs +/- EGF stimulation determined by immunoblotting. TCL: Total cell lysate; IAC: Integrin-associated complex enrichment sample. **C)** KEGG Pathway analysis of IAC dataset +/- EGF stimulation. Proteins were identified by LC-MS/MS and the fold change EGF/SFM (log_2_) calculated using ion intensity measurements. Proteins were submitted to DAVID and all significantly over-represented pathways are shown, with corresponding number of identified proteins representing each KEGG pathway term (Counts) and Benjamini corrected p-values. **D)** Representation of consensus adhesome and meta-adhesome dataset coverage. Node size corresponds to frequency of protein occurrence in IAC enrichment datasets analysed in Horton *et al* 2015. Inner ring (green fill/red border): consensus adhesome. Middle ring (green fill/grey border): meta-adhesome. Outer ring (grey node): not reported in the consensus or meta-adhesome. 68.3% of the consensus adhesome (41 proteins); 35.7% of the meta adhesome (836 proteins). Proteins in the outer ring were not reported in the consensus or meta adhesome (519 proteins). Total number of proteins = 1396. Nodes represent proteins and edges are known interactions. **E)** KEGG Pathway analysis of IAC enrichment +/- EGF stimulation. Proteins were identified by LC-MS/MS and the fold change EGF/SFM (log_2_) calculated using ion intensity measurements. Proteins were submitted to DAVID and four significant KEGG_Pathway terms are shown: Ribosome (no consistent change in enrichment +/-EGF) Focal adhesion (de-enriched following EGF), Regulation of actin cytoskeleton (de-enriched following EGF) and Endocytosis (de-enriched following EGF). Proteins of interest are highlighted with black arrows. N = 2, error bars = standard deviation (s.d.). Green dashed line represents 95% confidence interval.

Following sample validation, isolates were analysed by mass spectrometry. Total normalised ion abundance was quantified using Progenesis LC-MS. In both unstimulated and EGF-stimulated conditions, total ion abundance was comparable and the distribution of the mean normalised abundance was very similar (data not shown); demonstrating equal loading between both IAC samples and equal mass spectrometry performance. Consequently, any change observed was therefore due to the effect of EGF stimulation and not unequal total protein amount. Proteins were quantified using Progenesis LC-MS by alignment of the corresponding chromatograms and the alignment quality of in-gel trypsin digested samples was good (Score: 96.6%-98.5%) indicating that equivalent peptide ion peaks could be reliably compared. Therefore, downstream analyses were performed using relative quantification of EGF-induced fold changes generated using the ion intensity measurements.

To gain an initial overview of the data, all identified proteins were mapped onto the curated consensus- and meta-adhesome^17^ to determine the level of similarity, and disparity, between the new dataset and previously published IAC enrichment data (Figure 1D). Unlike previous IAC enrichment datasets, we used culture conditions that enable integrin-mediated engagement of a multiple ECM molecules. However, despite these differences, and the fact that IACs were isolated from epithelial cells, 68.3% of the consensus adhesome (41 proteins) was identified in the dataset. Moreover, 35.7% of the meta-adhesome (836 proteins) was detected. However, in addition to meta-adhesome components, a further 519 other proteins were isolated. This divergence could potentially be due to the complex nature of the ligand, enabling engagement of different integrin heterodimers, or due cellular context.

Identified proteins were subjected to KEGG pathway analysis (Figure 1C), Gene Ontology Enrichment Mapping (Figure S1A) and Gene Ontology clustering (Figure S1B/C). KEGG analysis analyses identified functional pathways that were enriched in the total dataset, including ‘Ribosome’, Proteasome’, ‘Focal adhesion’, ‘Regulation of actin cytoskeleton’ and ‘Endocytosis’ (Figure 1C). Gene Ontology Enrichment Mapping and clustering demonstrated that EGF stimulation induced a significant decrease in the abundance of adhesion and cytoskeletal proteins (Figure S1A-C). This change was closely coupled to a decrease in endocytic proteins (cluster 4) and suggested a potential mechanism of receptor internalisation and adhesion turnover.

To highlight EGF-induced changes to specific proteins which mapped to the KEGG Pathway terms ‘Focal adhesion’, ‘Regulation of actin cytoskeleton’ and ‘Endocytosis’, proteins were plotted against their mean fold change (Figure 1E). The majority of proteins within these terms demonstrated a mean fold change below the overall mean, indicating EGF-dependent de-enrichment (including integrins and EGFR). By contrast, extracellular proteins (including matrix proteins collagen, laminin and fibronectin and growth factors such as PDGFs) demonstrated very little change; suggesting that EGF-induced changes were specific to intracellular protein complexes. While adhesion complex, cytoskeletal and endocytic proteins consistently decreased following EGF stimulation, proteins that represented the ‘Ribosome’ KEGG pathway did not exhibit a consistent pattern of change, despite being highly enriched and over-represented in the dataset.

We next analysed network topology, in order to identify putative proteins with novel roles in integrin and EGFR crosstalk (Figure 2A). Proteins that were potential binding partners of both β1 integrin and EGFR, based on reported interactions with both receptors within the PPI network, were identified and termed "one-hop intersect" proteins (Figure 2A/B). The majority of one-hop intersect proteins were well-characterised adhesion proteins such as talin, FAK (Ptk2), α5 integrin, Src and Lyn. The membrane proteins CD44 (hyaluronan receptor), CD82 (tetraspannin) and CD98 (Slc3A2, amino acid transporter) were also identified as intersecting proteins. Shc1 is a well-known downstream effector of EGF signalling^18^, and PKCα (Prkca) is a serine/threonine protein kinase that regulates a number of cellular signalling cascades^19^. As with the previous KEGG and Gene Ontology analysis, network analysis demonstrated that 11/13 one-hop intersect proteins demonstrated a substantial decrease in abundance following EGF stimulation.

**Figure 2:**
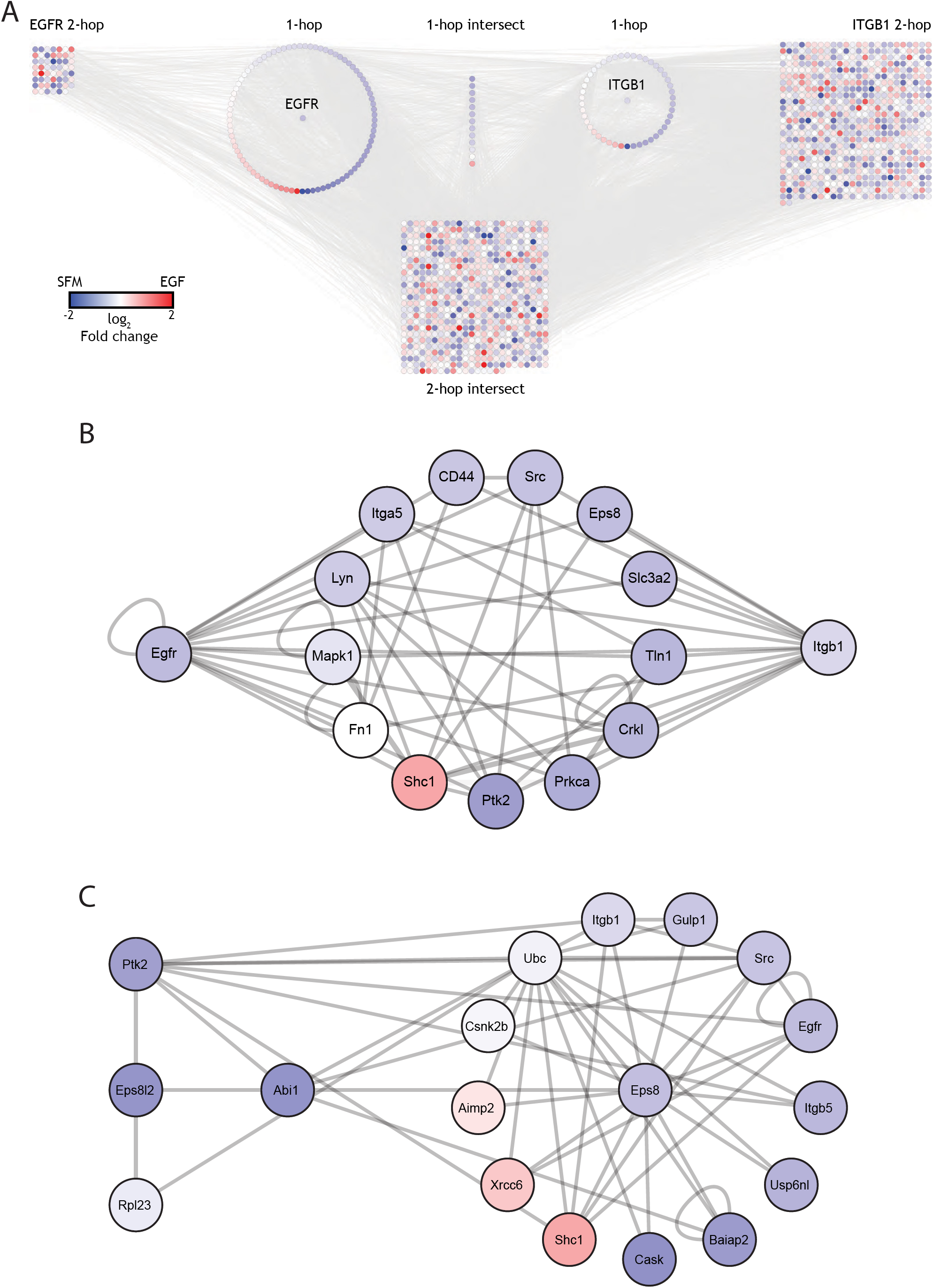
Network analysis of adhesion receptor-growth factor crosstalk identifies Eps8 as a key regulator of signal integration. **A)** Network analysis using ion intensity quantification of adhesion complex enrichment from EpH4 MECs +/- EGF. Ion intensities of proteins identified by LC-MS/MS were quantified using Progenesis LC-MS. Proteins were mapped onto protein-protein interaction network and shaded according to fold-change EGF/SFM (log_2_) following 5 minutes EGF stimulation. Proteins displayed: one- and two-hop neighbourhoods of β1 integrin (ITGB1) and EGF receptor (EGFR). **B)** Analysis of the β1 integrin-EGFR one-hop intersect sub-network of adhesion complexes. one-hop intersect sub-networks between ITGB1 and EGFR quantified using ion intensities. Shading indicates fold-change EGF/SFM (log_2_) following 5 minutes EGF stimulation. **C)** Analysis of the Eps8 and Eps8L2 one-hop sub-networks. one-hop sub-networks of Eps8 and Eps8L2 quantified using ion intensities. Shading indicates fold-change EGF/SFM (log_2_) following 5 minutes EGF stimulation.

Together, bioinformatic network and ontological analyses revealed a substantial decrease in the abundance of adhesion regulatory proteins and co-ordinators of endocytosis within 5 minutes of EGF stimulation. Given the overall decrease in proteins associated with Focal Adhesion, Regulation of Actin Cytoskeleton and Endocytosis KEGG terms, we reasoned that such a de-enrichment could be triggered by receptor internalisation and adhesion complex disassembly; a mechanism that would be consistent with EGF-induced receptor endocytosis and adhesion complex turnover. Therefore, we assessed whether any proteins identified within the one-hop intersect were associated with endocytic processes. While individual proteins within the one-hop intersect did not represent members of the Endocytosis Biological Process GO term or KEGG pathway, Eps8 was identified as a binding partner of RN-tre (Usp6nl) (Figure 2C). RN-tre is a GTPase-activating protein that forms a complex with Eps8 and functions as a negative regulator of Rab5 activity^20-24^.

Based on the EGF-dependent recruitment of Eps8 to adhesion sites, identification of endocytosis by GO and pathway analysis as a key EGF-dependent biological process at IACs, and the regulatory functions of, and binding partners associated with, Eps8, we identified Eps8 as putative regulator of EGFR and integrin crosstalk. Moreover, the inter-connectivity of Eps8 within the proteomic network indicated that Eps8 could have a functional role integrating growth factor mediated adhesion responses.

Interestingly, a related Eps8-family member, Eps8-like 2 (Eps8L2), was also identified in IACs and exhibited a similar level of de-enrichment following EGF stimulation. So Eps8L2 was also investigated further. However, it is notable that Eps8L2 was only identified in the two-hop intersect of α5β1 integrin and EGFR interactors and was not as inter-connected within the network as Eps8 (Figure 2C).

To validate the proteomic analyses, immunoblotting confirmed that Eps8 and Eps8L2 were enriched in isolated IACs and levels of detection were reduced following acute EGF stimulation. Moreover, de-enrichment of the consensus adhesome protein, paxillin, was also triggered by EGF, confirming the reduction identified by mass spectrometry (Figure 1E; Gene name PXN; Focal adhesion and Regulation of actin cytoskeleton KEGG pathways).

Having identified Eps8 as a putative regulatory protein in isolated IACs, indirect immunofluorescence was used to determine the subcellular distribution of Eps8. EpH4 and MEF cells were co-stained with Eps8, the canonical adhesion complex protein talin and phalloidin to assess Eps8 localisation to focal adhesions and actin, respectively. Fibroblasts were used in addition to EpH4 cells, as fibroblasts form robust load-bearing adhesion structures. Co-localisation between Eps8 and talin was observed in both EpH4 and MEF cell lines (Figure 3B/C), demonstrating that Eps8 is recruited to sites of cell-matrix interaction. However, while limited co-localisation was observed between Eps8 and Eps8L2, Eps8L2 was largely absent from talin-positive focal adhesion structures (Figure 3D). This observation was consistent with the reduced connectivity of Eps8L2 in isolated IACs and suggested that Eps8 may have more direct adhesion regulatory functions than Eps8L2.

**Figure 3:**
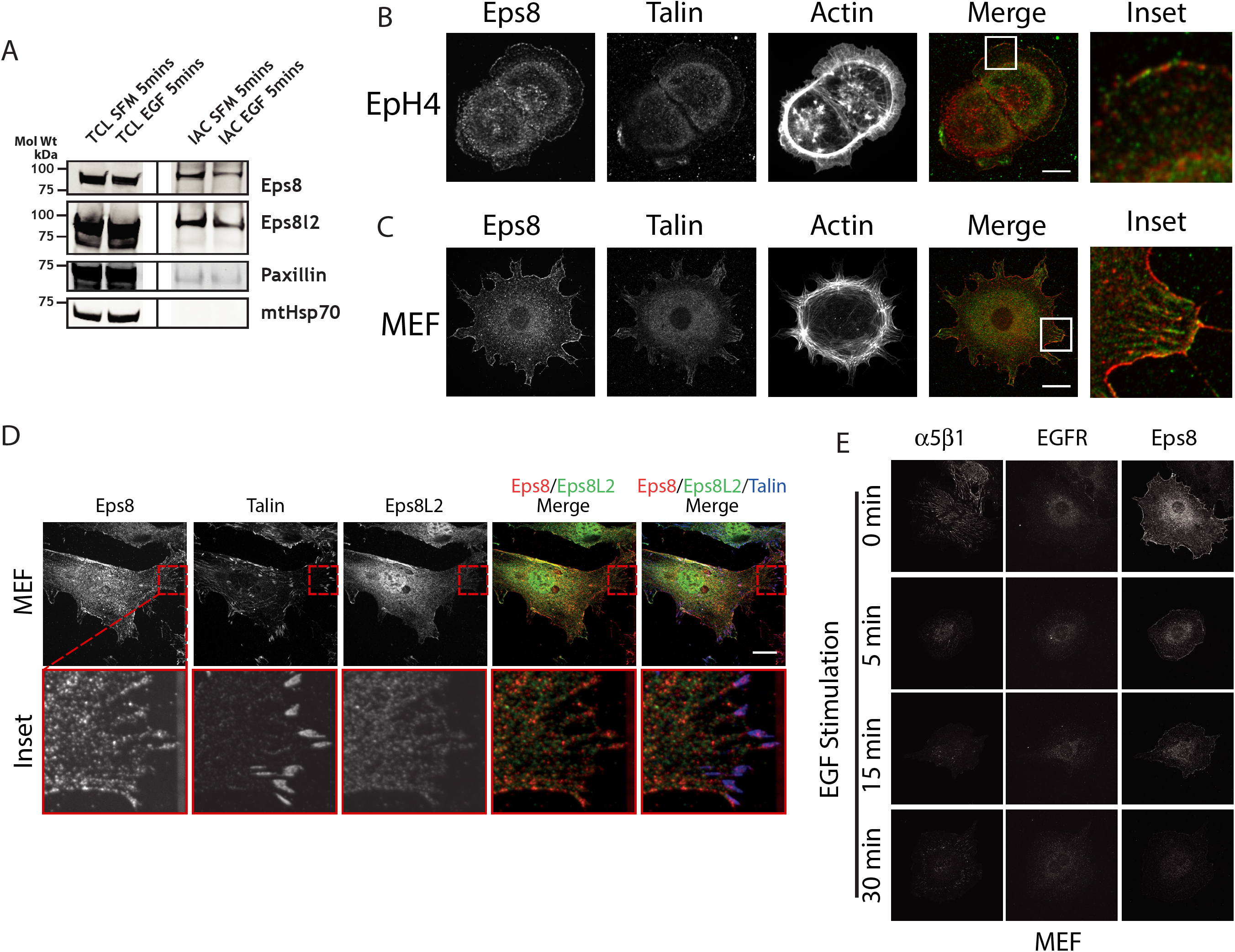
Eps8 is recruited to IACs and regulated by EGF stimulation. **A)** Validation of EGF-induced changes in IACs enriched from EpH4 MECs +/- EGF stimulation determined by immunoblotting. TCL: Total cell lysate; IAC: Integrin-associated complex enrichment sample. **B & C)** Eps8 localises to integrin-associated adhesion complexes. Immunofluorescence micrographs showing subcellular localisation of Eps8 (red) and talin (green) in **(B)** EpH4 cells (Scale bar = 10 μm) and **(C)** MEFs (Scale bar = 20). White boxes correspond to region of interest for insets. **D)** Eps8, but not Eps8L2, co-localises with talin at adhesion complexes in MEFs. Red boxes correspond to region of interest for insets. Lower panel: higher amplification image insets. Scale bar = 20 μm. All images are maximum z-slice projections and representative cells from three independent replicates. **E)** EGF stimulation promotes disassembly of α5β1 integrin-dependent adhesion complexes and redistribution of Eps8. Immunofluorescence micrographs showing α5β1, EGFR and Eps8 during a time-course of 10 ng/ml EGF stimulation in MEFs. Images are maximum z-slice projections and representative cells from three independent replicates.

Initial proteomic analyses were performed using conditions that enabled engagement of a mixed ECM microenvironment, including collagens and fibrillar ECM macromolecules that were synthesised and/or remodelled by the cells. However, the only integrin heterodimer identified within the one-hop intersect was the fibronectin-binding integrin, *α*5*β*1 (Figure 2B). Moreover, the only ECM molecule identified within the one-hop intersect was fibronectin. Importantly, while levels of the *α*5 and *β*1 integrin subunits in IACs were reduced by EGF stimulation, the levels of fibronectin (Fn1) were unchanged (Figure 2B); suggesting that EGF stimulation did not reduce *α*5*β*1 levels in IACs by modulating the availability of ligand. So we sought to determine whether EGF stimulation regulated the subcellular distribution of *α*5*β*1 and Eps8. Consistent with the ontological and network analyses (Figures 1/S1/2), EGF stimulation induced a rapid (with <5mins) loss of *α*5*β*1-containing adhesion complexes and redistribution of Eps8 from the cell periphery and adhesion sites (Figures 3E/S2A). This coincided with endocytosis of EGFR, but it is notable that *α*5*β*1 and EGFR did not appear to co-accumulate in endocytic vesicles. Interestingly, reformation of *α*5*β*1-dependent adhesion structures was observed 30-60 mins post-stimulation.

### Eps8 constrains EGFR and α5β1 integrin internalisation

Ontological analysis suggested that endocytosis was a key biological process differentially regulated by EGF stimulation at IACs. While the direct effect of Eps8 on integrin internalisation has never been determined, Eps8 forms a complex with RN-tre that inhibits Rab5 and suppresses EGFR endocytosis ^23^. The detection of both Eps8 and RN-tre in isolated IACs, and the role that RN-tre plays in Rab5-dependent heterodimer-specific integrin endocytosis ^25^ led us to consider whether Eps8 and Eps8L2 regulate integrin and/or EGFR trafficking. The EGF-dependent de-enrichment of α5β1 in IACs, and the relationship between α5β1, Eps8 and RN-tre in the proteomic networks suggested that the complex may have a role in α5β1 endocytosis.

To determine the effect of Eps8 on EGFR and integrin endocytosis, constitutive and EGF-stimulated internalisation rates of EGFR and integrin α5β1 were assessed by biochemical endocytosis assays in control, Eps8 and Eps8L2 knockdown MEFs (Ctrl KD, Eps8 KD and Eps8L2 KD, respectively). Under unstimulated conditions, Ctrl KD cells exhibited low levels of constitutive endocytosis of both α5β1 and EGFR (Figure 4A). Stimulation with EGF increased the endocytosis of both α5β1 and EGFR; indicating a degree of co-regulation between the two receptors. However, Eps8 KD cells exhibited high levels of endocytosis of both α5β1 and EGFR, even in the absence of in the absence of EGF-stimulation. Thus, siRNA-mediated Eps8 knockdown increased constitutive endocytosis of both α5β1 and EGFR, to levels comparable to EGF stimulation of control cells. Indeed, stimulation with EGF in Eps8 KD cells slightly increased the rate of α5β1 and EGFR endocytosis relative to control knockdown (Figure 4A). By contrast, Eps8L2 knockdown did not significantly affect either constitutive or ligand stimulated endocytosis of α5β1 or EGFR (Figure 4B). Together these data show that Eps8 regulates endocytosis of both α5β1 and EGFR and suggest that Eps8 functions to limit or constrain receptor internalisation in the absence of EGF stimulation. Consequently, EGF stimulation relieves Eps8-mediated inhibition of α5β1 and EGFR endocytosis. However, consistent with its distinct subcellular localisation (Figure 3D), Eps8L2 does not directly contribute to α5β1 and EGFR endocytosis.

**Figure 4:**
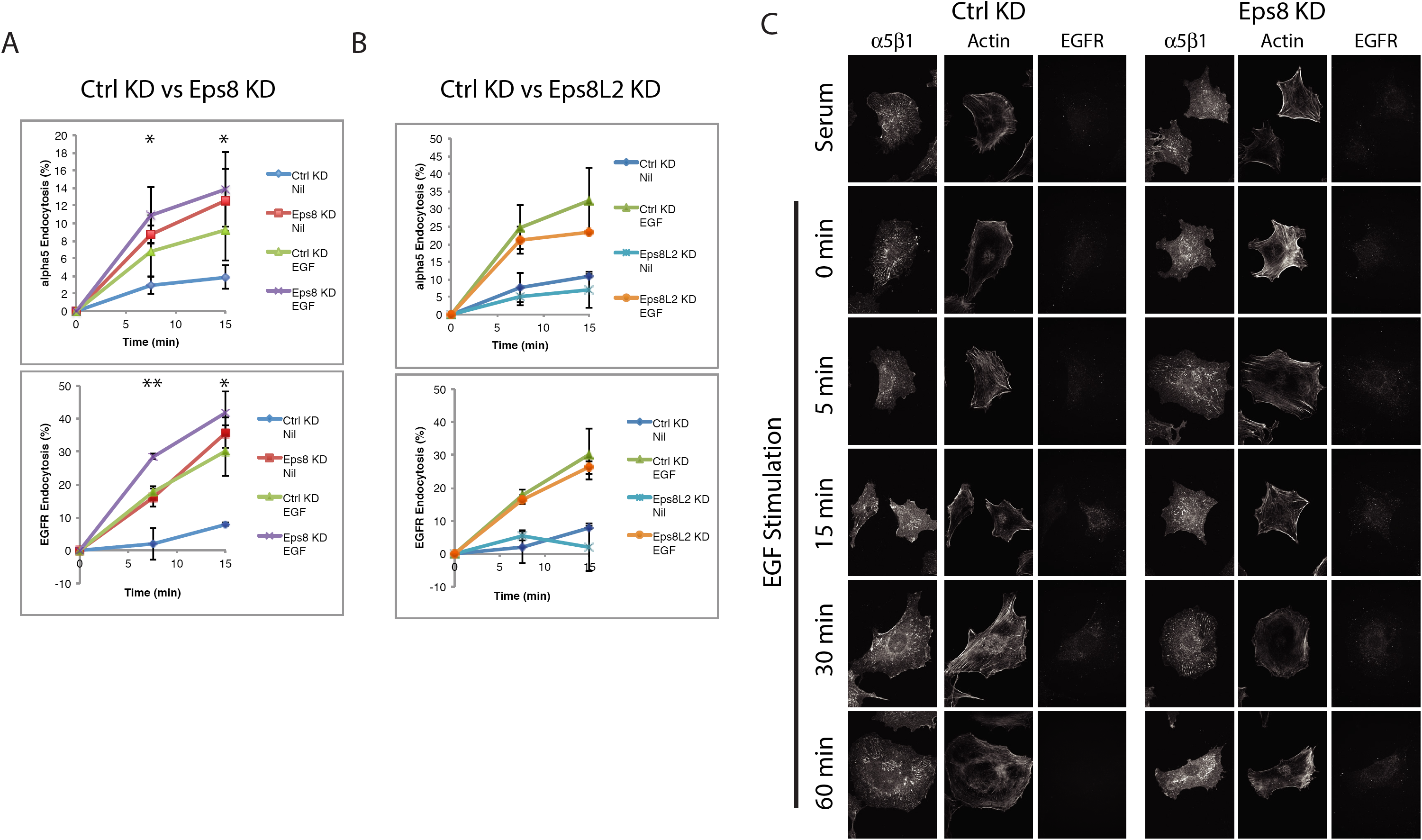
Eps8 constrains α5β1 and EGFR endocytosis in the absence of EGF stimulation. **A&B)** Endocytosis of integrin α5β1 and EGFR under unstimulated constitutive (serum-starved) and EGF stimulated (10ng/ml EGF) conditions. EGF Stimulation increases endocytosis of both α5β1 and EGFR. **A)** Ctrl KD vs Eps8 KD MEFs; Eps8 KD increases unstimulated constitutive and EGF stimulated endocytosis of α5β1 and EGFR. **B)** Ctrl KD vs Eps8 KD MEFs; Eps8L2 KD does not dysregulate α5β1 or EGFR endocytosis. α5β1 endocytosis N=7, EGFR endocytosis N=5. Duplicate plates per condition. Data points plotted as means with SEM error bars. Statistical test = two-tailed t-test assuming unequal variance. Black asterisks: Constitutive endocytosis rates in Eps8 KD compared with Ctrl KD cells (* = p< 0.05; ** = p< 0.01). **C)** Eps8 regulates adhesion complex disassembly. Eps8 is required for maintenance of adhesion complex organisation, and EGF-dependent adhesion complex disassembly. MEFs were serum starved then stimulated with 10 ng/ml EGF. Images are maximum z-slice projections. Representative cells from three replicates.

### Eps8 regulates EGF-mediated adhesion disassembly and Rab5 activity

Adhesion disassembly is closely linked to endocytosis, as dissociation of integrin-associated cytoskeletal components is required to permit formation of endocytic complexes and internalisation of integrins and associated proteins from the cell membrane ^26, 27^. As Eps8 restricts integrin α5β1 and EGFR internalisation, we assessed whether Eps8 has a role in regulating the disassembly or remodelling of adhesions. Adhesion complex organisation was therefore assessed by immunofluorescence under basal and EGF-stimulated conditions with Eps8 knockdown (Figure 4C/S2B). Under steady-state serum-containing conditions, α5β1 distribution was similar in control and Eps8 knockdown cells. However, while following serum-starvation α5β1 was organised in large adhesion complexes in control cells, Eps8 KD cells formed small disorganised α5β1-containing adhesion complexes (Figure 4C/S2B). As observed previously in untransfected cells (Figure 3E/S2A), EGF stimulation in control knockdown cells, caused rapid disassembly of α5β1-dependent adhesion complexes, prior to re-assembly that is initiated from 30 minutes. However, siRNA-mediated knockdown of Eps8 prevented disassembly of α5β1 adhesions in response to acute EGF stimulation and the IACs remained disorganised (Figure 4C/S2B). Together these data indicate that Eps8 is required for the organisation and EGF-mediated disassembly of adhesion complexes, which is likely related to Eps8-mediated regulation of α5β1 endocytosis.

Rab5 is a trafficking regulatory small GTPase that plays key roles in receptor endocytosis and endosomal maturation. Rab5 has been implicated in both EGFR and integrin endocytosis^28-30^. Rab5 has been reported to associate directly with integrins and promotes focal adhesion disassembly^30, 31^. Given that Eps8 regulates EGF-dependent adhesion disassembly, integrin and EGFR endocytosis, and forms a complex with the RabGAP, RN-tre^21, 22^, we assessed whether EGF stimulation directly regulates Rab5 activity and examined the impact that suppressing Eps8 has on Rab5. Stimulation of control cells with EGF induced rapid and substantial activation of Rab5 (Figure 5A); providing a mechanism by which EGF can trigger α5β1 integrin and EGFR endocytosis and drive adhesion disassembly (Figure 4A/B/C &S2A). However, siRNA knockdown of Eps8 enhanced basal levels of Rab5 activity (Figure 5B). These data are consistent with the ability of Eps8 to form a complex with the negative regulator of Rab5, RN-tre. to repress Rab5 activity^23^. Eps8 has the ability to directly bind integrin β-subunits^32^. So, as RN-tre was identified with Eps8 in isolated IACs and followed a similar pattern of de-enrichment following EGF stimulation (Figure 2), it is possible that the integrin-binding capacity of Eps8 recruits RN-tre to IACs. Such a recruitment mechanism would enable spatial control of receptor trafficking mechanisms at sites of cell-matrix interaction; whereby Eps8 regulates localised RN-tre recruitment to sites of integrin-engagement, in order to limit Rab5 activity in the absence of a "disassembly signal" such as EGF stimulation. Such a mechanism suggests that IACs are primed for rapid endocytosis-mediated disassembly, but that this is prevented by local Eps8-dependent suppression of Rab5 activity.

**Figure 5:**
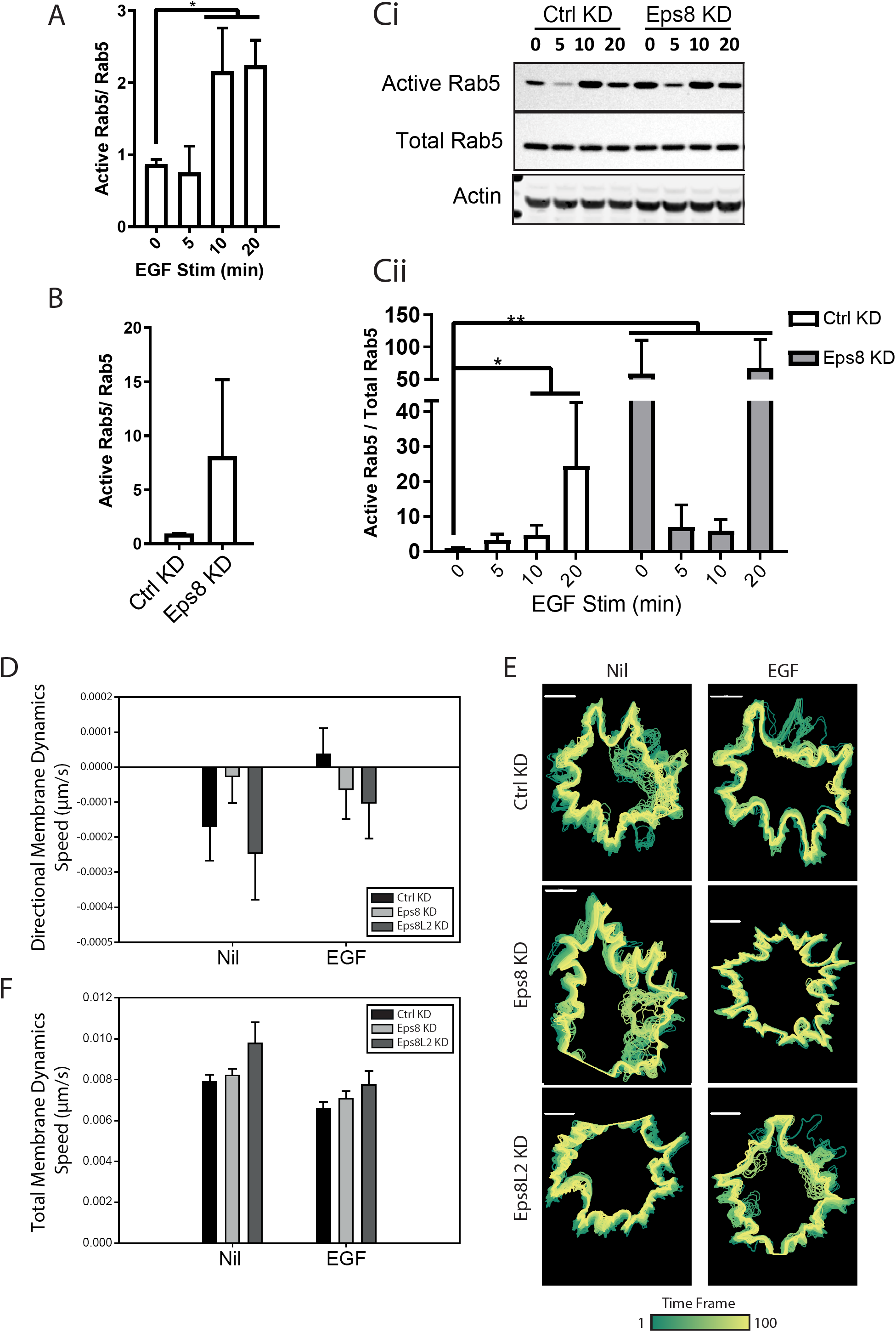
Eps8 regulates Rab5 activity and EGF stimulation relieves Eps8-dependent cellular contractility. **A)** EGF stimulation activates Rab5 activity. Rab5 activity (Rab5 GTP) assessed by effector pull-down assay. Time-course of 10 ng/ml EGF stimulation in serum-starved control MEFs. Quantification of active Rab5 (GST-R5BD-associated Rab5) relative to total Rab5 protein levels. N=3 error bars = SEM. **B)** Eps8 suppresses basal Rab5 activity. Rab5 activity (Rab5 GTP) assessed by effector pull-down assay. Basal levels of Rab5 activity in serum-starved Ctrl KD and Eps8 KD. Quantification of active Rab5 (GST-R5BD-associated Rab5) relative to total Rab5 protein levels. N=4, error bars = SEM. White bar = Ctrl KD, grey bar = Eps8 KD. **C)** Eps8 suppresses basal Rab5 activity. Rab5 activity (Rab5 GTP) assessed by effector pull-down assay. Time-course of 10 ng/ml EGF stimulation in serum-starved Ctrl KD and Eps8 KD. **Ci)** Shows representative immunoblotting. **Cii)** Quantification of active Rab5 (GST-R5BD-associated Rab5) relative to total Rab5 protein levels. N=5, error bars = SEM. White bar = Ctrl KD, grey bar = Eps8 KD. **D-F)** Effect of Eps8 KD on membrane protrusion and contractility dynamics. GFP-LifeAct-transfected MEFs were imaged on cell-derived matrices for one hour pre-and post-stimulation with 10 ng/ml EGF. Imaging was resumed approximately 10 minutes after EGF stimulation. Cell membrane tracking analysis was performed using the QuimP (V 18.02.01) plug-in for ImageJ and processed using MatLab function *readQanalysis.m*. **D)** Directional membrane dynamics speed (μm/s) for Ctrl KD (black), Eps8 KD (light grey) and Eps8L2 KD (dark grey) cells under serum-starved (Nil) and EGF stimulated conditions. Positive values indicate net membrane protrusion and negative values indicate net membrane contraction. N = 3; Error bars = SEM. **E)** Total membrane dynamics speed (μm/s) for control (black), Eps8 KD (light grey) and Eps8L2 KD (dark grey) cells under serum-starved (Nil) and EGF stimulated conditions. Values correspond to mean membrane speed (including protrusion and contraction), regardless of directionality. N = 3; Error bars = SEM. **F)** Cell tracks for representative single cells in which all cell outlines are overlaid and coloured according to frame number. Scale bar = 10 μm

It is important to note that, despite the fact that Eps8 KD cells exhibit high levels of basal Rab5 activity, EGF stimulation is able to regulate Rab5 activation in the absence of Eps8 (Figure 5C); inducing an initial suppression of Rab5 activity, followed by substantial Rab5 activation. Thus, while Eps8 serves to repress receptor endocytosis, this mechanism is de-coupled from the ability of EGF to directly stimulate Rab5 activity. This is consistent with our observation that, despite dysregulating EGFR endocytosis, Eps8 knockdown does not affect EGF-induced MAPK or Akt signalling (Figure S3). In normal physiology, functional de-coupling of mechanisms that restrain and trigger endocytosis would be essential to enable precise spatiotemporal control of receptor engagement.

### Role of Eps8 in EGF-dependent signalling and cytoskeletal dynamics

Eps8 has been linked to EGFR signalling, as overexpression of Eps8 increases cell proliferation in response to EGF^33^. As Eps8 regulates EGF-dependent Rab5 activity (Figure 5A-C) and EGFR/integrin endocytosis (Figure 4A), and therefore receptor bioavailability, we assessed whether Eps8 regulated ligand-dependent EGFR downstream signalling. Phosphorylation of canonical downstream signalling effectors was assessed in response to EGF. However, following siRNA-mediated knockdown of Eps8 or Eps8L2, no significant effect was observed on phosphorylation of EGFR or its downstream targets ERK or Akt in response to EGF (Figure S3A-C). Consistent with Lanzetti *et al*.^23^, these data indicate that Eps8 and Eps8L2 do not have a direct role in regulating EGF-dependent EGFR downstream signalling.

Eps8 can form a complex with Abi1 and Sos1 to regulate Rac1 activity and actin dynamics. Abi1 functions as a scaffold, linking Eps8 to the bifunctional Ras/Rac-GEF Sos1^23^. Eps8 association promotes Sos1 activity, and the tri-complex exhibits Rac-specific GEF activity^34^. Therefore, we examined whether Eps8 regulated basal or EGF-dependent Rac1 activity. However in MEFs, despite activating MAPK and Akt signalling, EGF stimulation did not induce transient activation of Rac1 activity and siRNA-mediated suppression of either Eps8 or Eps8L2 had no effect on basal or EGF-stimulated Rac1 activity (Figure S3D).

Together, these data indicate that Eps8 regulates EGF-dependent Rab5 activity and controls endocytosis of α5β1 and EGFR in MEFs. By contrast, Eps8 knockdown did not perturb either EGFR signalling or Rac1 activity. This is consistent with the notion that the Eps8-Abi1-Sos1 complex is regulated by RTK downstream signalling. PI3K is recruited to active RTKs and activated by Ras, to produce PIP3, which in turn activates Rac-GEFs including Sos1 ^35^.

We have identified a key role for Eps8 in the regulation of α5β1 and EGFR endocytosis and EGF-dependent adhesion turnover. Coordinated integrin endocytosis and recycling are required for spatiotemporal control of integrin-mediated adhesions and cytoskeletal dynamics, which is essential for membrane protrusion and retraction during cell migration^36, 37^. We therefore assessed the role of Eps8 in EGF-dependent control of cytoskeletal dynamics. Live-cell imaging was used to quantitatively analyse membrane protrusion and retraction activity on cell-derived matrices. When considering membrane dynamics, both control and Eps8L2 knockdown cells exhibited net contractile activity under serum-starved conditions and EGF stimulation relieved this contractility activity (Figure 5D/E). However, Eps8 knockdown inhibited cellular contractility in unstimulated cells and no additional effect of EGF stimulation was observed (Figure 5D/E). No changes were observed in the non-directional membrane motility data (Figure 5F), demonstrating overall dynamics were unchanged and only directionality (i.e. protrusion vs contraction) was influenced. Thus, Eps8 is required to sustain membrane contractility in the absence of EGF and for EGF-dependent release of cellular contractility.

## SUMMARY

In this study we employed a novel proteomic approach to dissect adhesion receptor-GFR crosstalk and identified a key regulatory mechanism controlling α5β1 integrin and EGFR functions. Analysis of adhesion signalling networks, following acute EGF stimulation, allowed a global and dynamic view of adhesion and growth factor receptor crosstalk to be assembled. Analysis of network topology identified Eps8 as a putative node integrating α5β1 integrin and EGFR functions:

Further analysis revealed:

- EGF stimulation promotes internalisation of both α5β1 and EGFR
- Eps8 constrains α5β1 and EGFR endocytosis in the absence of EGF stimulation
- Eps8 is required for maintenance of adhesion complex organisation and EGF-dependent adhesion complex disassembly
- EGF stimulation triggers Rab5 activity and Eps8 is required for EGF-dependent Rab5 activation
- Eps8 is required to sustain membrane contractility in the absence of EGF and for EGF-dependent release of cellular contractility

Together, these data suggest that Eps8 is recruited to integrin-associated adhesion complexes and serves to restrict unchecked α5β1 and EGFR endocytosis. As such, Eps8 retains integrin-mediated signalling complexes at the cell-matrix interface, yet has the capacity to respond rapidly to EGF stimulation to drive receptor internalisation, adhesion disassembly and reduce cellular contractility. We propose that during tissue morphogenesis and repair, Eps8 functions to spatially and temporally constrain endocytosis, and engagement, of α5β1 and EGFR in order to precisely co-ordinate adhesion disassembly, cytoskeletal dynamics and cell migration.

## MATERIALS AND METHODS

### Antibodies and reagents

All reagents were obtained from Sigma-Aldrich unless otherwise stated. Antibodies used for immunoblotting were mouse anti-Eps8 (BD Biosciences), rabbit anti-Eps8l2 (Proteintech Ltd.), rabbit anti-MAPK1 (Santa Cruz Biotechnology), rabbit anti-MAPK p44/42 T202/Y204 (Cell Signalling Technology), rabbit anti-ILK (Abcam), mouse anti-mitochondrial Hsp70 (Thermo Scientific Pierce), and mouse-anti Paxillin (BD Biosciences). Fluorophore-conjugated secondary antibodies for immunoblotting and immunofluorescence were from Life Technologies. Antibodies used for immunofluorescence were mouse anti-Eps8, mouse anti-Vinculin (Sigma-Aldrich), and Texas-Red Phalloidin (Life Technologies). Antibodies used for receptor internalisation assays were rat anti-α5 integrin (BD Biosciences), rat anti-β1 integrin (Millipore) and rabbit anti-EGFR (Genentech).

### Cell culture and transfection

EpH4 mouse mammary epithelial cells were routinely cultured in DMEM/F12 medium (Lonza) supplemented with 5% (v/v) FCS and 5 μg/ml Insulin. Immortalised mouse embryonic fibroblasts (IM-MEFs) were maintained in Dulbecco’s modified Eagle’s medium (DMEM) supplemented with 10% (v/v) FCS, 2 mM L-glutamine and 20 μg/ml recombinant mouse interferon-γ. Mouse Eps8 (#11: ACGACUUUGUGGCGAGGAA) and Eps8l2 (#10: UCGACUAUCUGUACGACAU) were silenced using ON-TARGETplus siRNA (Dharmacon, Thermo Fisher Scientific), 1.6μg per transfection. ON-TARGETplus Non-targeting siRNA was used as a negative control. Electroporation was performed using the Cell Line Nucleofector^®^ Kit V and Nucleofector 2b device (Lonza) according to the manufacturers’ instructions. A second round of transfection was performed after 48 hours.

### Adhesion complex isolation

Adhesion complexes were isolated from EpH4 cells according to Jones *et al.*^38^, with some modifications. Briefly, tissue culture plates were coated with 10 μg/ml collagen-I from rat tail. Prior to cell seeding, plates were washed with PBS, and blocked with 1% heat-denatured BSA (w/v from 99% stock) for 45 minutes-1 hour at RT. EpH4 cells seeded at 2-3×10^4^ cells/cm^2^ in DMEM/F12 containing 0.2% BSA, 5 ng/ml EGF and 5 μg/ml insulin for 16-18 hours. This prolonged period of incubation enabled the cells to synthesise and remodel a complex extracellular matrix. Cells were growth factor starved for a minimum of 4 hours in DMEM/F12.

Cells were treated with 5ng/ml EGF or left untreated (serum-free medium, SFM) for 5 minutes at 37°C/5% CO2. Media was removed and adhesions isolated directly with hypotonic water pressure held ∼1cm from the surface of the dish for 10-15 seconds per plate. Plates were washed immediately with ice-cold PBS and dried on ice for 1 minute. Excess liquid was removed and isolated adhesions were scraped into reducing sample buffer (RSB). Samples were denatured by heating to 95°C for 5 minutes. Proteins were precipitated in ice-cold acetone on dry-ice for a minimum of 3 hours. Precipitated samples were centrifuged at 16000g for 15 minutes at 4°C and washed three times with cold acetone. The resulting protein pellet was dried at room temperature (RT) and resuspended in the desired volume of RSB by shaking at 1400rpm for 30 minutes at 70°C followed by reduction for 5 minutes at 95°C. To load one SDS-PAGE gel, complexes were isolated from two 10cm dishes and resuspended in 35μl RSB prior to analysis by immunoblotting and mass spectrometry.

### SDS-PAGE and Immunoblotting

Cells were washed twice with PBS on ice and scraped into ice-cold lysis buffer (1% (v/v) TX-100, 150 mM NaCl, 50 mM Tris, 10 mM MgCl2, 5 mM EGTA pH 7.4, 5 mM EDTA pH 8.0, 0.5 mM AEBSF, 25 μg/ml aprotinin, 25 μg/ml leupeptin and 2 mM Na_3_VO_4_). Cell lysates were pre-cleared by centrifugation at 21000g for 10 minutes at 4°C. The supernatant was collected and resolved by SDS-PAGE. Proteins were transferred in transfer buffer (25 mM Tris-HCl, 190 mM glycine, 0.01% (w/v) SDS, 20% (v/v) methanol) at 30V for 90 minutes. Nitrocellulose membranes were blocked with blocking buffer diluted in PBS for 1 hour at RT. Membranes were probed with primary antibodies resuspended to the appropriate concentration in blocking buffer diluted in Tris Buffered Saline (TBS: 50 mM Tris, 150 mM NaCl pH 7.4) containing 0.01% (v/v) Tween-20 (TBST) and incubated for 1 hour at RT or overnight at 4°C. Membranes were washed with TBST and incubated with the appropriate fluorescent dye-conjugated secondary antibody diluted in TBST for 45 minutes at RT. Membranes were washed with TBST and visualised using the Odyssey IR Imaging System (700nm and 800nm channels, 169μm resolution without focus offset).

### Mass spectrometry analysis

Protein samples were resolved by SDS-PAGE and gel lanes were sliced prior to in-gel tryptic digestion. Peptide samples were analysed by liquid chromatography-tandem mass spectrometry (LC-MS/MS) using an UltiMate^®^ 3000 Rapid Separation LC (RSLC, Dionex Corporation) coupled to an Orbitrap Elite mass spectrometer (Thermo Fisher Scientific). Peptide mixtures were separated using a 250 mm x 75μm i.d. 1.7 μM BEH C18, analytical column (Waters) using a 45-min linear gradient from 1 to 25% (vol/vol) acetonitrile in 0.1% (vol/vol) formic acid at a flow rate of 200 nl/min. Peptides were selected for fragmentation automatically by data dependant analysis. RAW data files were searched using Mascot Daemon (version 2.2.2, Matrix Science, London, UK), against a FASTA format protein sequence database with taxonomy of *Mus musculus* selected. Database searching was performed using an in-house Mascot server (http://msct.smith.man.ac.uk/mascot/home.html; Matrix Science, London, UK)^39^. For all database searches, carbamidomethylation of cysteine (+57.02 Da) was set as a fixed modification, and oxidation of methionine (-64.00 Da; +16.00 Da) allowed as a variable modification. Monoisotopic precursor mass values were used and only doubly and triply charged precursor ions were considered. Mass tolerances for precursor and fragment ions were 5ppm and 0.5 Da respectively. Alternatively, ion intensity analysis was performed from RAW files using Progenesis LC-MS (version 4.1, Non-linear dynamics, Newcastle-upon-Tyne, UK). Protein-protein interaction network analysis was performed using Cytoscape (versions 2.8.3 and 3.60) ^40, 41^. Proteins were mapped onto a human interaction network composed of the Protein Interaction Network Analysis (PINA, 10th December 2012) ^42, 43^, the Matrisome Project ^44-47^ and the Adhesome^16^. Gene Ontology analyses were performed using the Database for Annotation, Visualization and Integrated Discovery (DAVID, version 6.7).

### Receptor Internalisation Assays

Internalisation assays were performed as described previously^48^. High binding 96-well plates (Corning) were coated with 5 μg/ml antibodies in 0.05 M Na_2_CO_3_ pH 9.6 at 4°C overnight. Antibody coated plates were blocked with 5% (w/v) BSA/0.1% (w/v) Tween-20 in PBS at RT for 1 hour. IM-MEFs were allowed to spread onto 10cm dishes overnight and growth factor-starved in DMEM for a minimum of 4 hours. Cells were washed twice with ice-cold PBS on ice, and labelled with 0.13 mg/ml EZLink-Sulfo-NHS-SS-Biotin (Thermo Fischer Scientific) for 30 minutes at 4°C on ice under gentle rocking. Following labelling, excess biotin was removed with two washes of ice-cold KREBS buffer (118 mM NaCl, 25 mM NaHCO_3_, 4.8 mM KCl, 1.2 mM KH_2_PO_4_, 1.2 mM MgSO_4_, 11 mM glucose, 1.5 mM CaCl_2_.2H_2_0, 1.5 mM sodium pyruvate). To assess total biotin binding, 1 or 2 plates were maintained in KREBS buffer on ice until lysis. In addition, control plates were maintained in KREBS buffer on ice until the cell surface reduction step. For the remaining plates, internalisation was permitted by adding prewarmed DMEM (37°C), with and without the presence of 10 ng/ml EGF, and transferring the plates to a 37°C incubator for the indicated times. Following internalisation, medium was aspirated and cells were washed with ice-cold KREBS buffer on ice. Control and assay plates were washed with pH 8.6 Buffer (50 mM Tris-HCl pH 7.5, 100 mM NaCl pH 8.6). Cell surface reduction was achieved by addition of 3.75 mg/ml sodium 2-mercaptoethanesulfonate (MesNa) in pH 8.6 Buffer for 30 minutes on ice with gentle rocking. The cell surface reduction label was removed and an additional 30-minute reduction step performed. All plates were washed twice with KREBS buffer, drained and lysed in 100-250μl lysis buffer (200 mM NaCl, 75 mM Tris-HCl, 15 mM NaF, 1.5 mM Na_3_VO_4_, 7.5 mM EDTA, 7.5 mM EGTA, 1.5% (w/v) Triton X-100, 0.75% (w/v) Igepal CA-630, 50 μg/ml leupeptin, 50 μg/ml aprotinin and 1 mM AEBSF) per plate. Lysates were clarified by centrifugation at 21000g for 10 minutes at 4°C. ELISA plates were washed twice with PBS/0.1% Tween and thoroughly drained before addition of 50μl lysate to the appropriate wells. Binding was permitted overnight (16-18 hours) at 4°C. Unbound material was removed by 4 washes with PBS/0.1% Tween and the plate was drained thoroughly. Extravadin^®^ Peroxidase (1 mg/ml in PBS-Tween/1% BSA) was added to each well (50μl/well) for 1 hour at 4°C. Following incubation, wells were washed 4 times with PBS-/0.1% Tween and drained. Development substrate (40 mM 2,2**′**-Azinobis(3-ethylbenzothiazoline-6-sulphonic acid) diammonium salt (ABTS) in ABTS buffer (0.1 M Na Acetate, 0.05 M NaH_2_PO_4_ pH 5.0) and 2.5 mM H_2_O_2_) was added to each well and the absorbance read at 405nm at regular intervals prior to saturation of the signal. Percentage internalisation was expressed as a fraction of the total (non-reduced) cell surface label minus the background (reduced but not internalised) label.

### Indirect immunofluorescence

Cells were fixed with 4% (w/v) paraformaldehyde (PBS (-), pH 6.9) for 20 minutes at room temperature, then washed three times in PBS (-). Cells were then permeabilized for 3–4 minutes with 0.1% (v/v) Triton X-100 at room temperature followed by three 0.1/0.1 buffer (PBS + 0.1% BSA, 0.1% sodium azide) washes. Primary antibody incubations were for 45 minutes at room temperature, followed by three washes in 0.1/0.1 buffer. Samples were incubated with secondary antibody, with or without phalloidin (1:400 in 0.1/0.1 buffer), for 45 minutes at room temperature protected from light. Samples were then washed twice in PBS (-) and once in Milli-Q water, before mounting with Prolong Gold anti-fade mountant (Molecular Probes Invitrogen) on glass Superfrost^®^ Plus glass slides (Thermo Scientific).

Samples were imaged using the Zeiss 3i Marianas spinning disk confocal system using a 63x/1.4 oil objective. Downstream image processing was performed using Image J FIJI.

### Rac1 activity assays

Rac1 activity was evaluated using a GST-PAK1-PBD (p21 binding domain) fusion protein beads. 24h after a second round of transfection with either All-star negative control, siEps8 or siEps8L2 MEFs were treated with EGF (10ng/ml) for 0, 5, 10, 20 min and lysed with ice cold lysis buffer (25mM Tris×HCl, pH 7.2, 150mM NaCl, 5mM MgCl_2_, 1% NP-40, 2.5% glycerol, 50?mM NaF, 1?mM Na_3_VO_4_, 0.5 mM AEBSF 2ug/ml aprotinin and 2ug/ml leupeptin). Lysates were clarified by centrifugation at 16,000 g, 4°C, for 15 min. 800?ug protein was incubated with GST-PAK1-PBD-immobilised beads with gentle rotation for 1h at 4°C. The beads were washed three times with lysis buffer followed by elution of bound proteins with 2x reducing sample buffer (125mM Tris×HCl, pH 6.8, 2% glycerol, 4% SDS (w/v) and 0.05% bromophenol blue, 5% β-mercaptoethanol). Solubilised proteins were resolved by SDS-PAGE and levels of active Rac1 and total Rac1 proteins were analysed by immunoblotting.

### Rab5 activity assays

MEFs were seeded at 8.9 x 10^4^ cells/cm^2^ and incubated overnight in full growth medium. Cells were serum-starved for 4 hours prior to stimulation with 10 ng/ml EGF for 0, 5, 10 and 20 mins. Cells were lysed in lysis buffer containing 25 mM Tris-HCl pH 7.2, 150 mM NaCl, 5 mM MgCl_2_, 1% NP40, 5 % glycerol and protease inhibitors by scraping. Lysates were clarified by centrifugation at 16000xg for 15 min. Lysate supernatants were incubated with glutathione-sepharose beads pre-coated with 30ug of GST-RAB5BD during 30 min at 4°C in a rotating shaker followed by 3 washes with the lysis/binding/wash buffer. Finally, samples were boiled in Laemmli buffer and analysed by immunoblotting. The GST-RAB5BD construct, comprising GST-conjugated to the Rab5-binding domain of rabaptin-5, was a generous gift from Dr Vicente Torres from Faculty of Odontology of University of Chile.

### Generation of Cell-derived Matrices

Cell-derived matrices were produced by stimulating TIF fibroblasts to produce extracellular matrix, and subsequently removing the cellular material leaving the intact structure^49^. Fibroblasts were cultured over time on pre-treated 13 mm coverslips (VWR) in a 24 well plate format.

Coverslips were washed three times in PBS (-) then incubated with 0.2% sterile gelatin (from porcine skin) for one hour at 37°C. Coverslips were again washed in PBS (-), cross linked with 1% (v/v) sterile glutaralydehyde in PBS (-) for 30 minutes at room temperature. After another PBS (-) wash, the crosslinker was then quenched with 1 M sterile glycine in PBS (-) for 20 minutes at room temperature. The coverslips were then washed with PBS (-) and left to equilibrate in the appropriate culture medium (DMEM, 10% (v/v) FCS) at 37°C, prior to cell seeding.

TIFs were detached with 1 x trypsin-EDTA as described and seeded at a concentration of 5 x 10^4^ cells per well on the pre-treated coverslips. Cells were then cultured overnight at 37°C 8% CO2. If cells were sub-confluent at this stage, they were cultured longer until they reached confluence. The medium on confluent cells was changed to the same medium, supplemented with 50 μg/ml ascorbic acid. The ascorbic acid supplemented media was replaced every other day, until denudation. Ascorbic acid stimulates collagen production and stabilises the matrix. Cells were cultured at 37°C, 8% CO_2_ for 8 days from the first ascorbic acid stimulation, prior to denudation.

On the day of denudation, medium was removed and cells were washed with PBS (-), before adding pre-warmed (37°C) extraction buffer (20 mM NH4OH, 0.5% (v/v) Triton X-100 in PBS (-)). Cells were lysed for 2 minutes, by which point no intact cells were visible. Extraction buffer was removed, and cells washed in PBS (+). Residual DNA was digested with 10 μg/ml DNase I (Roche) in PBS (+) for 30 minutes at 37°C, 5% CO2. The DNase solution was removed, followed by two PBS (+) washes. At this point the cell-derived matrices were ready for use or stored at 4°C in PBS (+) supplemented with 1 x antibiotic antimycotic solution (100 x stock: 10,000 units penicillin, 10 mg streptomycin, 25 μg/ml amphotericin B, Sigma), for up to a month.

Prior to use, cell-derived matrices were washed twice in PBS (+) before blocking in heat-denatured BSA for 30 minutes at room temperature. Matrices were then washed in PBS (+) and equilibrated in the appropriate medium for the assay.

### Cytoskeletal Dynamics Analysis

MEFs were transfected with control, Eps8 or Eps8L2 siRNA. The second transfection was a co-transfection, containing 4 μg of LifeAct-GFP (Thistle Scientific) in addition to the siRNA oligonucleotides. LifeAct stains filamentous (F-actin) without interfering with actin dynamics^50^. The next day cell-derived matrices were washed in PBS (-), then blocked in heat denatured BSA for 30 minutes at room temperature. Matrices were then washed again in PBS (-), then in phenol red free reduced serum Opti-MEM™ (Fisher Scientific) and left to equilibrate at 37°C. Transfected MEFs were seeded at a density of 5 x 10^3^ per compartment in a 35mm diameter 4 compartment CELLview™ glass bottom cell culture dish (Greiner Bio-One).

16 hours later the media in each compartment was replenished, and cells were imaged using a Zeiss 3i Spinning disk confocal microscope Marianas™ SDC live cell imaging system, using a 63x/1.4 aperture oil objective. Images were captured at 40 second intervals. After an hour of imaging, cells were stimulated with EGF (10 ng/ml), and then imaged after a brief interval (2-3 minutes) required for capture focusing, for another hour.

Cellular protrusive activity and motility was quantified using the QuimP software set of plug-ins for ImageJ (Till Bretschneider, University of Warwick)^51^.

## ACKNOWLEDGEMENTS

This work was supported by a Cancer Research UK PhD studentship (NP), Wellcome Trust PhD studentship 102379/Z/13/Z (JRT), Breast Cancer Now Grant 2015MayPR507 (HM), North West Cancer Research grants CR1010 & CR1143 (KIW), Wellcome Trust grant 092015 (EJK/JDH/AB/MJH), BBSRC Biotechnology and Biological Sciences Research Council studentship from the Doctoral Training Programme (AG).

The mass spectrometers and microscopes used in this study were purchased with grants from BBSRC, Wellcome Trust, University of Liverpool (Institute of Translational Medicine) Strategic Fund and University of Manchester Strategic Fund.

## SUPPLEMENTARY FIGURE LEGENDS

**Figure S1:**
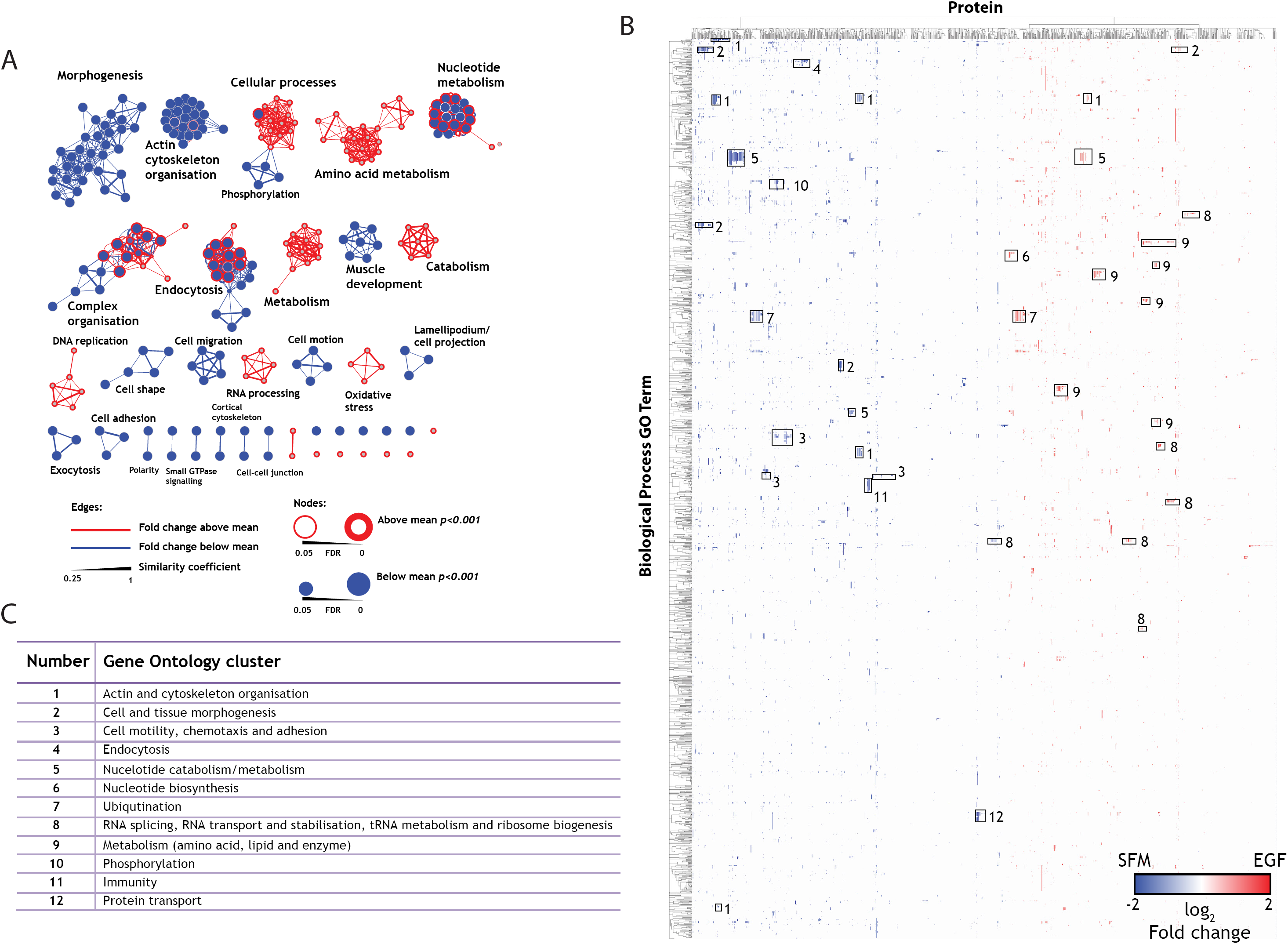
Gene ontology analysis of adhesion receptor-growth factor crosstalk. **A)** Gene Ontology Enrichment Map analysis of IACs +/- EGF. Proteins identified by LC-MS/MS and quantified using ion intensity measurements were submitted to DAVID for Gene Ontology analysis (Biological_Process_ALL). All proteins were subdivided either above or below the mean fold change EGF/SFM (log_2_) in normalised abundance. Gene Ontology terms were displayed using Enrichment Map and clustered. Edge weight (similarity coefficient) indicates the overlap between GO terms. Node size corresponds to enrichment of data below the mean (FDR). Node border width corresponds to enrichment of data above the mean (FDR). p-value cut-off = 0.001, False Discovery Rate (FDR) cut-off = 0.05, Jaccard Coefficient = 0.25. **B)** Gene Ontology hierarchical clustering analysis of IACs +/- EGF. Proteins identified by LC-MS/MS and quantified using ion intensity measurements were submitted to GO miner Gene Ontology analysis. Biological Process GO terms and proteins were hierarchically clustered using an uncentered Pearson correlation (complete linkage). Proteins were shaded according to fold-change EGF/SFM (log_2_). Major Biological Process clusters are numbered. **C)** Major Biological Process clusters identified by Gene Ontology hierarchical clustering (Fig S1B); including: (1) Actin and cytoskeleton organisation, (2) Cell and tissue morphogenesis, (3) Cell motility, chemotaxis and adhesion, (4) Endocytosis and (5) Nucleotide catabolism/metabolism. Biological Process clusters that were identified as uniquely decreasing following EGF stimulation were clusters 3, 4, 10, 11, 12. Clusters that were only increasing following EGF stimulation were 6, 9 and 12.

**Figure S2:**
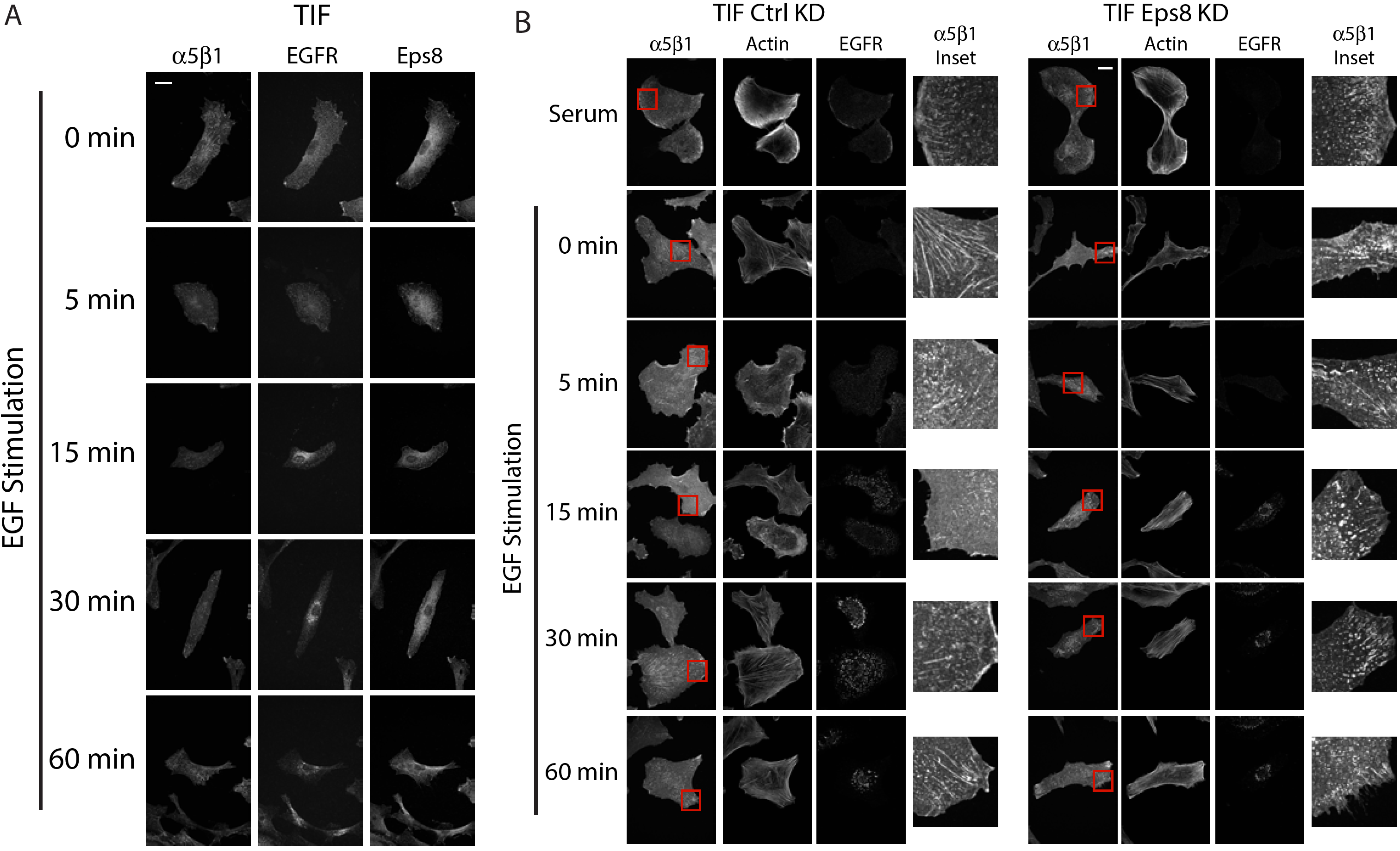
Eps8 suppresses EGF-induced α5β1 IAC disassembly. **A)** EGF stimulation promotes disassembly of α5β1 integrin-dependent adhesion complexes and redistribution of Eps8. Immunofluorescence micrographs showing α5β1, EGFR and Eps8 during a time-course of 10 ng/ml EGF stimulation in TIFs. Images are maximum z-slice projections and representative cells from three independent replicates. **B)** Eps8 regulates adhesion complex disassembly. Eps8 is required for maintenance of adhesion complex organisation, and EGF-dependent adhesion complex disassembly. TIF cells were serum starved and\ stimulated with 10 ng/ml EGF. Images are maximum z-slice projections. Representative cells from three replicates. Scale bars = 30 μm

**Figure S3:**
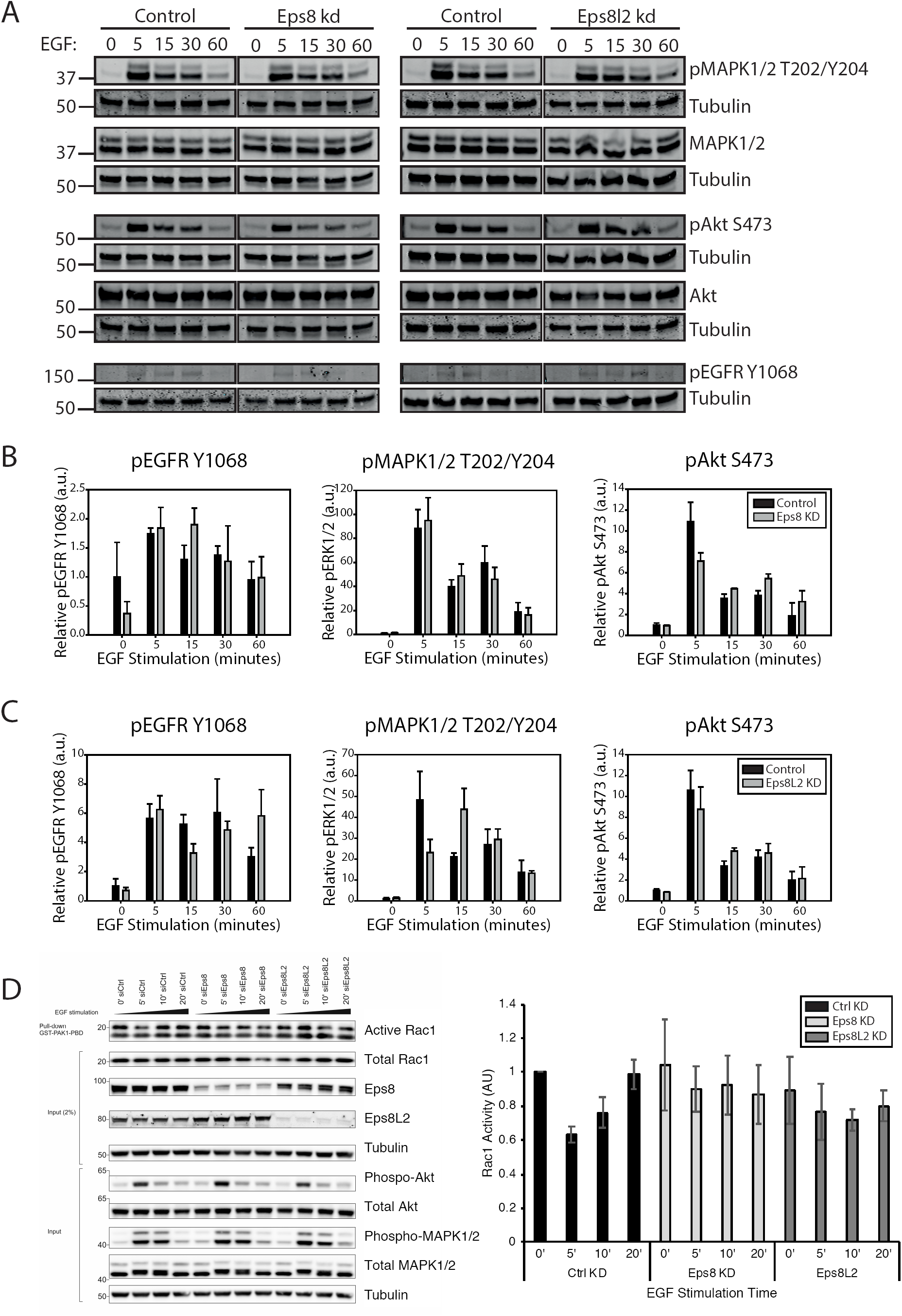
Eps8 knockdown does not modulate EGFR signalling. **A)** Time-course of 10 ng/ml EGF stimulation in serum-starved MEFs (Ctrl KD, Eps8 KD and Eps8L2 KD). EGF-dependent signalling assessed by Immunoblot analysis of phosphorylated EGFR, ERK1/2, Akt. pEGFR Y1068 normalised to tubulin, pMAPK1/2 T202/Y204 normalised to total MAPK and pAkt S473 normalised to total Akt. **B&C)** Quantification of phosphorylation relative to total protein levels or tubulin normalised to unstimulated control in Eps8 KD **(B)** or Eps8L2 KD cells **(C)**. N=3, error bars = SEM. Black bar = control siRNA, grey bar = Eps8 or Eps8L2 KD conditions (**B** and **C**, respectively). **D)** Rac1 activity (Rac1 GTP) assessed by effector pull-down assay. Time-course of 10 ng/ml EGF stimulation in serum-starved Ctrl KD, Eps8 KD and Eps8L2 KD MEFs. Quantification of active Rac1 (GST-PAK-associated Rac1) relative to total Rac1 protein levels. N=4, error bars = SEM. Black bar = control siRNA, light grey bar = Eps8, light grey bar = Eps8L2 KD.

## Abbreviations

ECM: Extracellular matrix
EGF: Epidermal growth factor
EGFR: Epidermal growth factor receptor
Eps8: Epidermal growth factor receptor pathway substrate 8
GFR: Growth factor receptor
IAC: Integrin-associated complex
MAPK: Mitogen activated protein kinase
MEC: Mammary epithelial cell
MEF: Mouse embryonic fibroblast
PKCα: Protein kinase Cα
TIF: T-antigen immortalised fibroblast

## References

1. Humphries, J.D., Paul, N.R., Humphries, M.J. & Morgan, M.R. Emerging properties of adhesion complexes: what are they and what do they do? Trends Cell Biol 25, 388–397 (2015).

2. Streuli, C.H. & Akhtar, N. Signal co-operation between integrins and other receptor systems. Biochem J 418, 491–506 (2009).

3. Reynolds, A.R. et al. Stimulation of tumor growth and angiogenesis by low concentrations of RGD-mimetic integrin inhibitors. Nat Med 15, 392–400 (2009).

4. Raab-Westphal, S., Marshall, J.F. & Goodman, S.L. Integrins as Therapeutic Targets: Successes and Cancers. Cancers (Basel) 9 (2017).

5. DuFort, C.C., Paszek, M.J. & Weaver, V.M. Balancing forces: architectural control of mechanotransduction. Nat Rev Mol Cell Biol 12, 308–319 (2011).

6. Ivaska, J. & Heino, J. Cooperation Between Integrins and Growth Factor Receptors in Signaling and Endocytosis. Annual review of cell and developmental biology (2011).

7. Horton, E.R. et al. Definition of a consensus integrin adhesome and its dynamics during adhesion complex assembly and disassembly. Nat Cell Biol 17, 1577–1587 (2015).

8. Horton, E.R. et al. Modulation of FAK and Src adhesion signaling occurs independently of adhesion complex composition. J Cell Biol 212, 349–364 (2016).

9. Robertson, J. et al. Defining the phospho-adhesome: phosphoproteomic analysis of integrin signalling. Nature Communications, In press (2015).

10. Byron, A. et al. A proteomic approach reveals integrin activation state-dependent control of microtubule cortical targeting. Nat Commun 6, 6135 (2015).

11. Schiller, H.B., Friedel, C.C., Boulegue, C. & Fassler, R. Quantitative proteomics of the integrin adhesome show a myosin II-dependent recruitment of LIM domain proteins. EMBO reports 12, 259–266 (2011).

12. Schiller, H.B. et al. beta1-and alphav-class integrins cooperate to regulate myosin II during rigidity sensing of fibronectin-based microenvironments. Nat Cell Biol 15, 625– 636 (2013).

13. Kuo, J.C., Han, X., Hsiao, C.T., Yates, J.R., 3rd & Waterman, C.M. Analysis of the myosin-II-responsive focal adhesion proteome reveals a role for beta-Pix in negative regulation of focal adhesion maturation. Nature cell biology 13, 383–393 (2011).

14. Winograd-Katz, S.E., Fässler, R., Geiger, B. & Legate, K.R. The integrin adhesome: from genes and proteins to human disease. Nature Reviews Molecular Cell Biology 15, 273–288 (2014).

15. Geiger, T. & Zaidel-Bar, R. Opening the floodgates: proteomics and the integrin adhesome. Current opinion in cell biology 24, 562–568 (2012).

16. Zaidel-Bar, R., Itzkovitz, S., Ma’ayan, A., Iyengar, R. & Geiger, B. Functional atlas of the integrin adhesome. Nature Cell Biology 9, 858–867 (2007).

17. Horton, E.R. et al. Definition of a consensus integrin adhesome and its dynamics during adhesion complex assembly and disassembly. Nat Cell Biol 17, 1577–1587 (2015).

18. Rozakis-Adcock, M. et al. Association of the Shc and Grb2/Sem5 SH2-containing proteins is implicated in activation of the Ras pathway by tyrosine kinases. Nature 360, 689–692 (1992).

19. Rosse, C. et al. PKC and the control of localized signal dynamics. Nat Rev Mol Cell Biol 11, 103–112 (2010).

20. Haas, A.K., Fuchs, E., Kopajtich, R. & Barr, F.A. A GTPase-activating protein controls Rab5 function in endocytic trafficking. Nat Cell Biol 7, 887–893 (2005).

21. Lanzetti, L. et al. Regulation of the Rab5 GTPase-activating protein RN-tre by the dual specificity phosphatase Cdc14A in human cells. J Biol Chem 282, 15258–15270 (2007).

22. Lanzetti, L., Palamidessi, A., Areces, L., Scita, G. & Di Fiore, P.P. Rab5 is a signalling GTPase involved in actin remodelling by receptor tyrosine kinases. Nature 429, 309– 314 (2004).

23. Lanzetti, L. et al. The Eps8 protein coordinates EGF receptor signalling through Rac and trafficking through Rab5. Nature 408, 374–377 (2000).

24. Matoskova, B., Wong, W.T., Nomura, N., Robbins, K.C. & Di Fiore, P.P. RN-tre specifically binds to the SH3 domain of eps8 with high affinity and confers growth advantage to NIH3T3 upon carboxy-terminal truncation. Oncogene 12, 2679–2688 (1996).

25. Palamidessi, A. et al. The GTPase-activating protein RN-tre controls focal adhesion turnover and cell migration. Curr Biol 23, 2355–2364 (2013).

26. Ezratty, E.J., Bertaux, C., Marcantonio, E.E. & Gundersen, G.G. Clathrin mediates integrin endocytosis for focal adhesion disassembly in migrating cells. The Journal of Cell Biology 187, 733–747 (2009).

27. Ezratty, E.J., Partridge, M.A. & Gundersen, G.G. Microtubule-induced focal adhesion disassembly is mediated by dynamin and focal adhesion kinase. Nat Cell Biol 7, 581– 590 (2005).

28. Barbieri, M.A. et al. Epidermal growth factor and membrane trafficking: EGF receptor activation of endocytosis requires Rab5a. J Cell Biol 151, 539–550 (2000).

29. Pellinen, T. & Ivaska, J. Integrin traffic. J Cell Sci 119, 3723–3731 (2006).

30. Pellinen, T. et al. Small GTPase Rab21 regulates cell adhesion and controls endosomal traffic of beta1-integrins. The Journal of Cell Biology 173, 767–780 (2006).

31. Mendoza, P. et al. Rab5 activation promotes focal adhesion disassembly, migration and invasiveness in tumor cells. Journal of cell science 126, 3835–3847 (2013).

32. Calderwood, D.A. et al. Integrin beta cytoplasmic domain interactions with phosphotyrosine-binding domains: a structural prototype for diversity in integrin signaling. Proc Natl Acad Sci U S A 100, 2272–2277 (2003).

33. Fazioli, F. et al. Eps8, a substrate for the epidermal growth factor receptor kinase, enhances EGF-dependent mitogenic signals. EMBO J 12, 3799–3808 (1993).

34. Scita, G. et al. EPS8 and E3B1 transduce signals from Ras to Rac. Nature 401, 290– 293 (1999).

35. Vanhaesebroeck, B. et al. Synthesis and function of 3-phosphorylated inositol lipids. Annu Rev Biochem 70, 535–602 (2001).

36. Paul, N.R. et al. alpha5beta1 integrin recycling promotes Arp2/3-independent cancer cell invasion via the formin FHOD3. J Cell Biol 210, 1013–1031 (2015).

37. Paul, N.R., Jacquemet, G. & Caswell, P.T. Endocytic Trafficking of Integrins in Cell Migration. Curr Biol 25, R1092–1105 (2015).

38. Jones, M.C. et al. Isolation of integrin-based adhesion complexes. Curr Protoc Cell Biol 66, 9 8 1–15 (2015).

39. Perkins, D.N., Pappin, D.J., Creasy, D.M. & Cottrell, J.S. Probability-based protein identification by searching sequence databases using mass spectrometry data. Electrophoresis 20, 3551–3567 (1999).

40. Demchak, B. et al. Cytoscape: the network visualization tool for GenomeSpace workflows. F1000Res 3, 151 (2014).

41. Smoot, M.E., Ono, K., Ruscheinski, J., Wang, P.L. & Ideker, T. Cytoscape 2.8: new features for data integration and network visualization. Bioinformatics 27, 431–432 (2011).

42. Cowley, M.J. et al. PINA v2.0: mining interactome modules. Nucleic Acids Res 40, D862–865 (2012).

43. Wu, J. et al. Integrated network analysis platform for protein-protein interactions. Nat Methods 6, 75–77 (2009).

44. Hynes, R.O. & Naba, A. Overview of the matrisome--an inventory of extracellular matrix constituents and functions. Cold Spring Harb Perspect Biol 4, a004903 (2012).

45. Naba, A. et al. The extracellular matrix: Tools and insights for the “omics” era. Matrix Biol 49, 10–24 (2016).

46. Naba, A. et al. The matrisome: in silico definition and in vivo characterization by proteomics of normal and tumor extracellular matrices. Mol Cell Proteomics 11, M111 014647 (2012).

47. Naba, A., Hoersch, S. & Hynes, R.O. Towards definition of an ECM parts list: an advance on GO categories. Matrix Biol 31, 371–372 (2012).

48. Roberts, M., Barry, S., Woods, A., van der Sluijs, P. & Norman, J. PDGF-regulated rab4-dependent recycling of alphavbeta3 integrin from early endosomes is necessary for cell adhesion and spreading. Curr Biol 11, 1392–1402 (2001).

49. Beacham, D.A., Amatangelo, M.D. & Cukierman, E. Preparation of extracellular matrices produced by cultured and primary fibroblasts. Curr Protoc Cell Biol Chapter 10, Unit 10 19 (2007).

50. Riedl, J. et al. Lifeact: a versatile marker to visualize F-actin. Nat Methods 5, 605–607 (2008).

51. Dormann, D., Libotte, T., Weijer, C.J. & Bretschneider, T. Simultaneous quantification of cell motility and protein-membrane-association using active contours. Cell Motil Cytoskeleton 52, 221–230 (2002).

